# BiocMAP: A Bioconductor-friendly, GPU-Accelerated Pipeline for Bisulfite-Sequencing Data

**DOI:** 10.1101/2022.04.20.488947

**Authors:** Nicholas J Eagles, Richard Wilton, Andrew E. Jaffe, Leonardo Collado-Torres

## Abstract

**Background:** Bisulfite sequencing is a powerful tool for profiling genomic methylation, an epigenetic modification critical in the understanding of cancer, psychiatric disorders, and many other conditions. Raw data generated by whole genome bisulfite sequencing (WGBS) requires several computational steps before it is ready for statistical analysis, and particular care is required to process data in a timely and memory-efficient manner. Alignment to a reference genome is one of the most computationally demanding steps in a WGBS workflow, taking several hours or even days with commonly used WGBS-specific alignment software. This naturally motivates the creation of computational workflows that can utilize GPU-based alignment software to greatly speed up the bottleneck step. In addition, WGBS produces raw data that is large and often unwieldy; a lack of memory-efficient representation of data by existing pipelines renders WGBS impractical or impossible to many researchers.

**Results:** We present BiocMAP, a Bioconductor-friendly Methylation Analysis Pipeline consisting of two modules, to address the above concerns. The first module performs computationally-intensive read alignment using *Arioc*, a GPU-accelerated short-read aligner. The extraction module extracts and merges DNA methylation proportions - the fractions of methylated cytosines across all cells in a sample at a given genomic site. Since GPUs are not always available on the same computing environments where traditional CPU-based analyses are convenient, BiocMAP is split into two modules, with just the alignment module requiring an available GPU. Bioconductor-based output objects in R utilize an on-disk data representation to drastically reduce required main memory and make WGBS projects computationally feasible to more researchers.

**Conclusions:** BiocMAP is implemented using Nextflow and available at http://research.libd.org/BiocMAP/. To enable reproducible analysis across a variety of typical computing environments, BiocMAP can be containerized with Docker or Singularity, and executed locally or with the SLURM or SGE scheduling engines. By providing Bioconductor objects, BiocMAP’s output can be integrated with powerful analytical open source software for analyzing methylation data.

## Background

The genome of many organisms is more than just a sequence of four nucleotides. These nucleotides can be chemically modified, and a common modification is the methylation of cytosines [1], which was discovered in mammals as early as DNA itself [2]. The percent of methylated cytosines was first measured across a significant portion of the human genome using methylation arrays [3]. With the advent of whole genome sequencing, whole genome bisulfite sequencing (WGBS) became a reality, allowing researchers to study methylated cytosines in different contexts (CpG, CpH) [4] and experimental settings [3]. However, population-scale studies have generally been limited to microarrays due to complexities in sample pre-processing required for WGBS. While methylation typically occurs primarily at cytosines in CpG context in almost all cell and tissue types, CpH-context methylation is present and plays a significant role in the brain [5].

Raw sequencing reads require several computational processing steps to produce DNA methylation proportions, a feature ready for statistical analysis. Among the most computationally demanding steps is alignment of reads to a reference genome, where software must consider alignments across reference sequences from 4 methylation states: methylated and unmethylated cytosines under directional and non-directional protocols. With currently available CPUs, alignment software such as *Bismark* [6] may require hours or days to align a single WGBS sample. The *Arioc* [7] aligner, which uses GPU acceleration to compute WGBS alignments, achieves processing speeds that are an order of magnitude faster without sacrificing accuracy or sensitivity. This provides a natural motivation for the implementation of a workflow that can use GPUs for alignment but CPUs for remaining processing steps. However, the use of GPUs for non-graphics-related tasks is still in its infancy, and GPU resources are sometimes not available on the same computing clusters where traditional CPU and memory resources are abundant.

We introduce BiocMAP [8], a **Bioc**onductor-friendly Nextflow-based [9] **M**ethylation **A**nalysis **P**ipeline for processing bisulfite-sequencing data into *bsseq* [10] R [11] objects that are readily analyzable with a number of Bioconductor [12] R packages. BiocMAP is split into two modules that can be executed in different computing environments; this can allow a researcher to align samples in a computing environment with ample GPU resources, but perform “methylation extraction” - calculating the fraction of methylated cytosines at a given genomic site - and remaining processing steps in an environment with more CPUs and memory.

## Results

### Overview

The BiocMAP workflow consists of a set of two modules - alignment and extraction, which together process raw WGBS reads in FASTQ format into Bioconductor-friendly [12] R [11] objects containing DNA methylation proportions essentially as a cytosine-by-sample matrix (**Figure 1**). In the first alignment module, an initial quality check is performed with *FastQC* [13], after which samples are trimmed with *Trim Galore!* [14], aligned to a reference genome with *Arioc* [7], and low-quality or duplicate mappings are filtered out.

**Figure 1.**
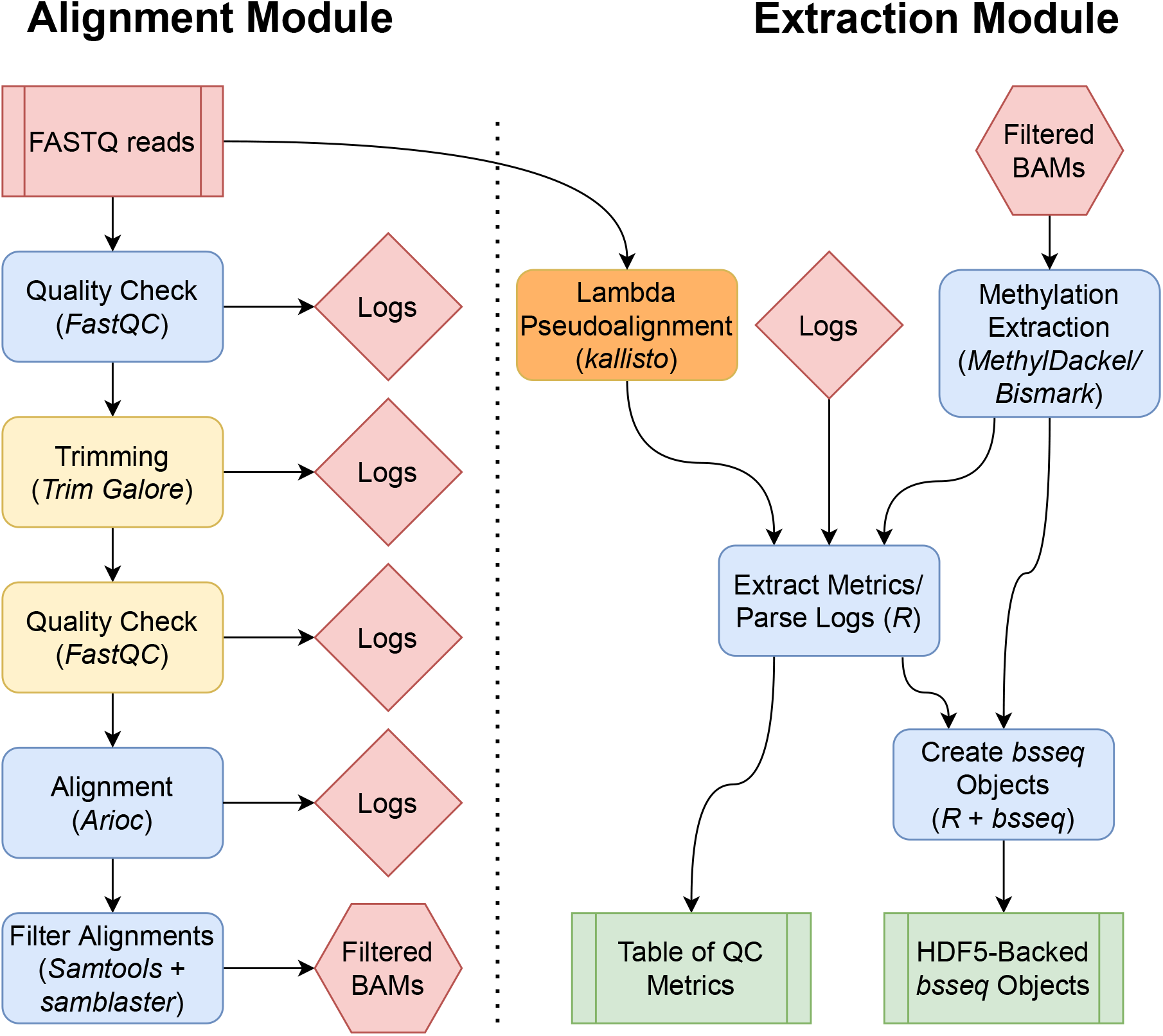
BiocMAP workflow overview. Diagram representing the conceptual workflow traversed by BiocMAP. The red box indicates the FASTQ files are inputs to the pipeline; green coloring denotes major output files from the pipeline; the remaining boxes represent computational steps. Yellow and orange-colored steps are optional or not always performed; for example, lambda pseudoalignment is an optional step intended for experiments with spike-ins of the lambda bacteriophage. Finally, blue-colored steps are ordinary processes which occur on every pipeline execution. Depending on the available high performance computing (HPC) systems, (a) both modules can be run sequentially on a HPC system with both GPUs and CPUs, or (b) the alignment module can be run on a GPU-powered HPC system, then files transferred to a CPU-based HPC system as well as updating file paths on the *rules.txt* file (dotted line), before running the extraction module on the CPU-based HPC system.

In the second extraction module, DNA methylation proportion extraction is performed within each sample using *MethylDackel* [15] or optionally *Bismark* [6], and the results are aggregated across samples into a pair of *bsseq* [10] R [11] objects for easy integration with a number of Bioconductor [12] packages to facilitate down-stream statistical analyses. The first *bsseq* object contains counts of methylated and unmethylated cytosines in CpG context across the entire reference genome, while the second object contains any additional cytosines in CpH context, when relevant (**Figure 2**). A summary table is also produced, compiling together metrics and statistics from trimming, alignment, and methylation extraction for each sample (Additional File 1: S1). Examples of information gathered include percent of reads concordantly aligned and percent of reads trimmed. This allows researchers to control for potential covariates and unintended sources of variation when performing downstream statistical analyses such as the identification of differentially methylated regions (DMRs) [10].

**Figure 2.**
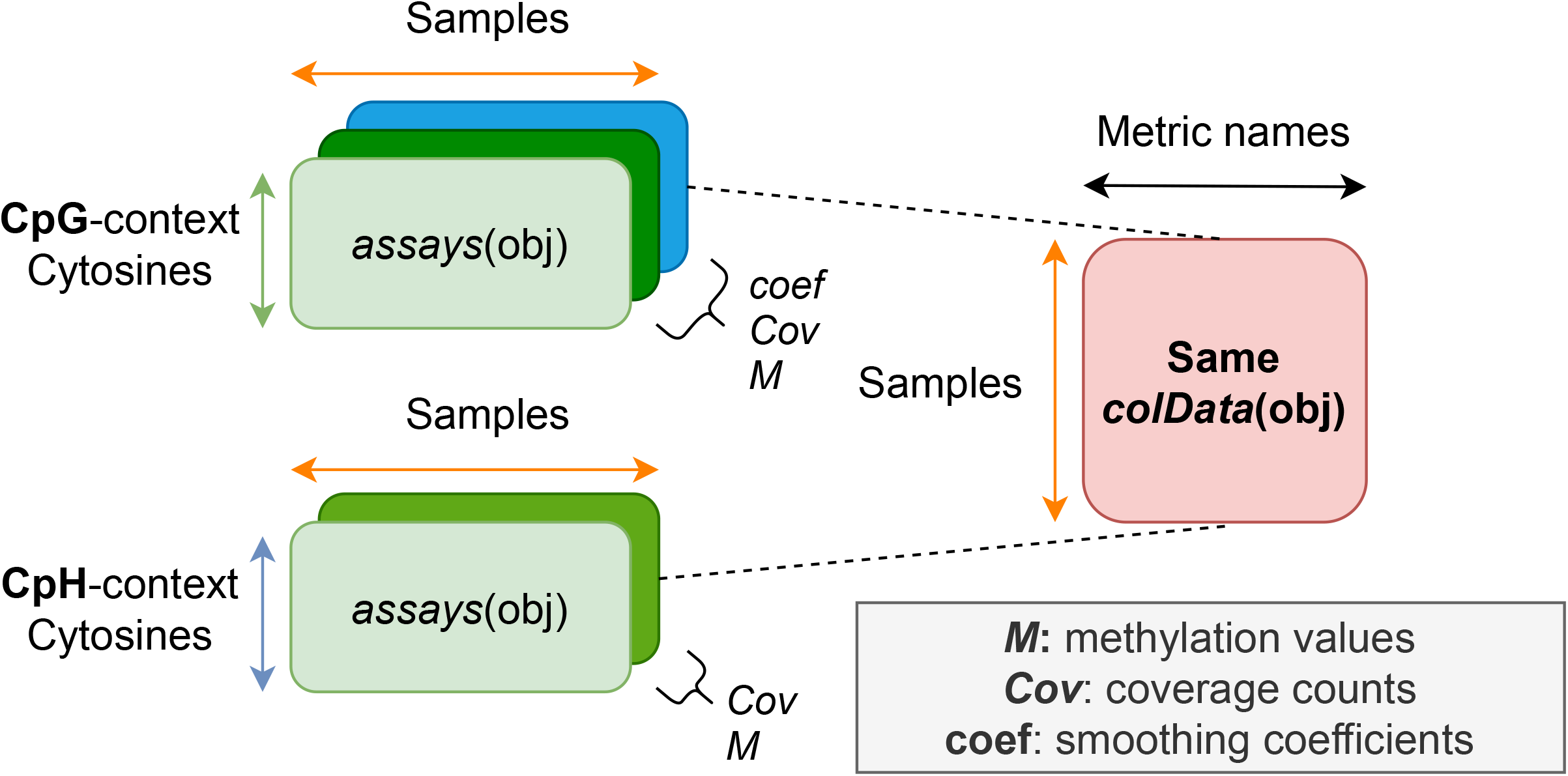
bsseq output objects. The major outputs from the extraction module are R objects from the *bsseq* Bioconductor package, which contain methylation proportion and coverage information at all cytosine loci in the human genome. *bsseq* extends the *SummarizedExperiment*class, which provides a general and popular format for storing genomics data and is memory efficient thanks to the *HDF5Array* backend. Two *bsseq* objects are produced, with one object containing cytosine sites in CpG context, and the other containing the remaining CpH loci.

### Application

To demonstrate how BiocMAP outputs might be used to perform statistical analysis and visualization on real WGBS data (Additional File 2), we used a publicly available dataset and illustrate an example analysis [8]. The dataset used includes 32 human brain dorsolateral prefrontal cortex (DLPFC) samples spanning developmental years from postnatal up to 23 years of age [16]. NeuN-based fluorescence-activated nuclear sorting was used to produce 8 glial and 24 neuronal samples from the homogenate DLPFC tissue. Prenatal samples present in Price et al. [5] were excluded for this analysis. Both modules of BiocMAP were executed on the data to produce output R objects that load into 23GB of memory. Execution times for each computational step in BiocMAP with this dataset were recorded and provide a guideline for other datasets (**Figure 3**). The execution time boxplots that can be generated as part of a Nextflow [9] “execution report”, produced by including the *-with-report* command-line option to the appropriate BiocMAP execution script (Additional File 1: S2).

**Figure 3.**
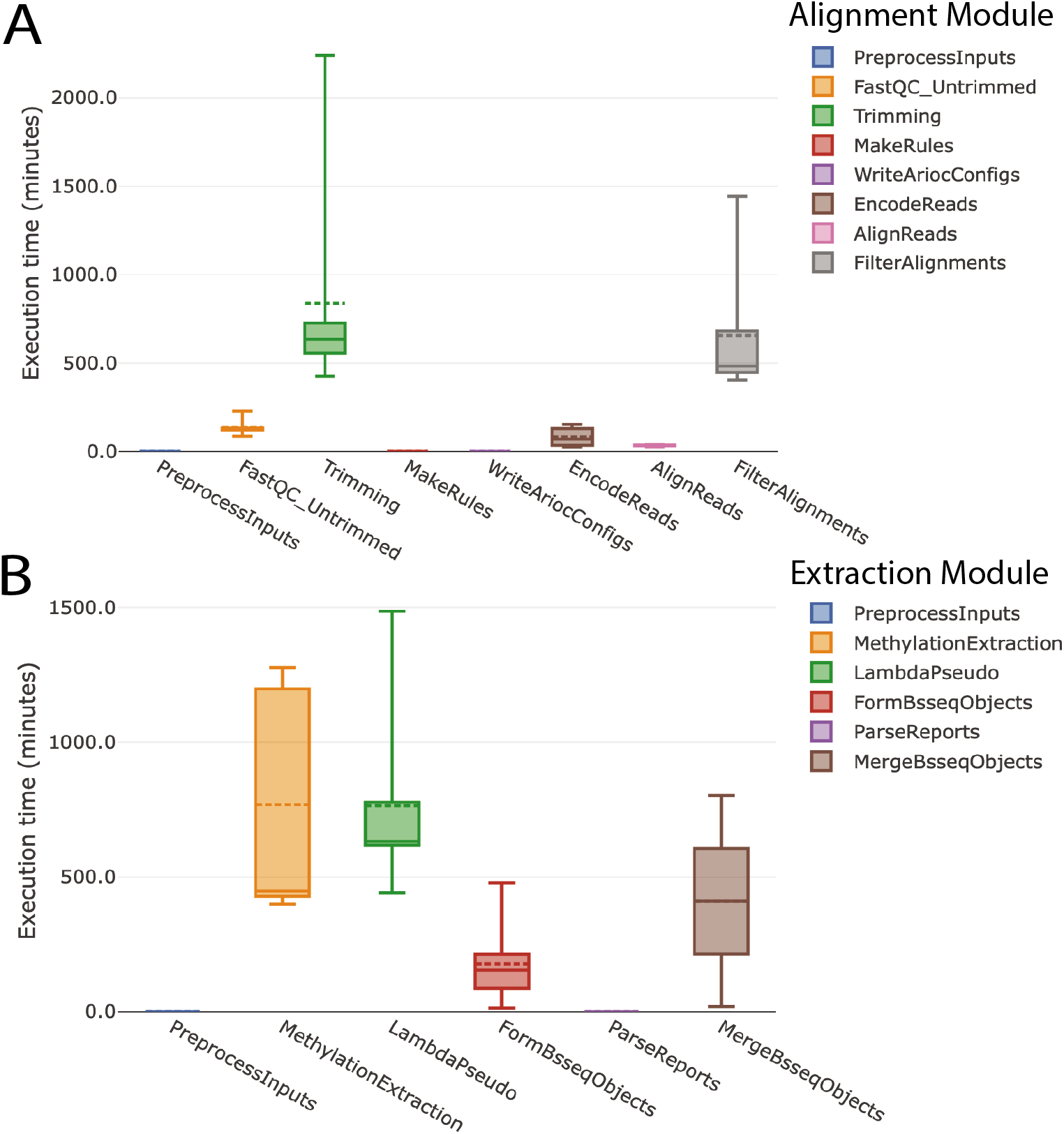
Process Run-times from Combined Execution Reports. Wall-clock run-times for each process in the alignment and extraction BiocMAP modules, (A) and (B) respectively, are plotted for the 32-sample subset of the Price et. al dataset [16]. Individual boxplots for each module, combined here for illustration purposes, are one several plot types included in an HTML execution report generated by including the *-with-report* command-line option to BiocMAP, as made possible through the Nextflow [9] framework. A given process, or computational step, in the BiocMAP workflow may be executed for one of more samples in the dataset; boxplots here summarize the distribution of run-times across all executions of each process type.

In the example analysis, we show how to attach external sample metadata to the *bsseq* [10] objects and produce several exploratory plots (Additional File 2). For example, we compared the estimated bisulfite-conversion rate (produced by BiocMAP) across neuronal and glial samples (provided by the dataset metadata; Additional File 2). We examined the relationship between trinucleotide methylation contexts (CpG, CHG, CHH), which shows higher methylation rates for neurons compared to glia as well as higher correlation between the methylation rate in CpG context against CHH or CHG context in neurons versus glia (**Figure 4A**). The proportion of cytosines with methylation higher than 10% is not significantly different under the CpG context, but is on the CpH context (CHH or CHG) when comparing neurons versus glia (**Figure 4B**). This proportion doesn’t change across ages 0 to 23. The original study describing this dataset identified differentially methylated regions (DMRs) between cell types [5]. By providing R/Bioconductor objects, BiocMAP’s outputs can easily be integrated with other R/Bioconductor packages that provide statistical and visualization methods. For example, DMRs can be visualized with *bsseq* [10] (**Figure 4C**).

**Figure 4.**
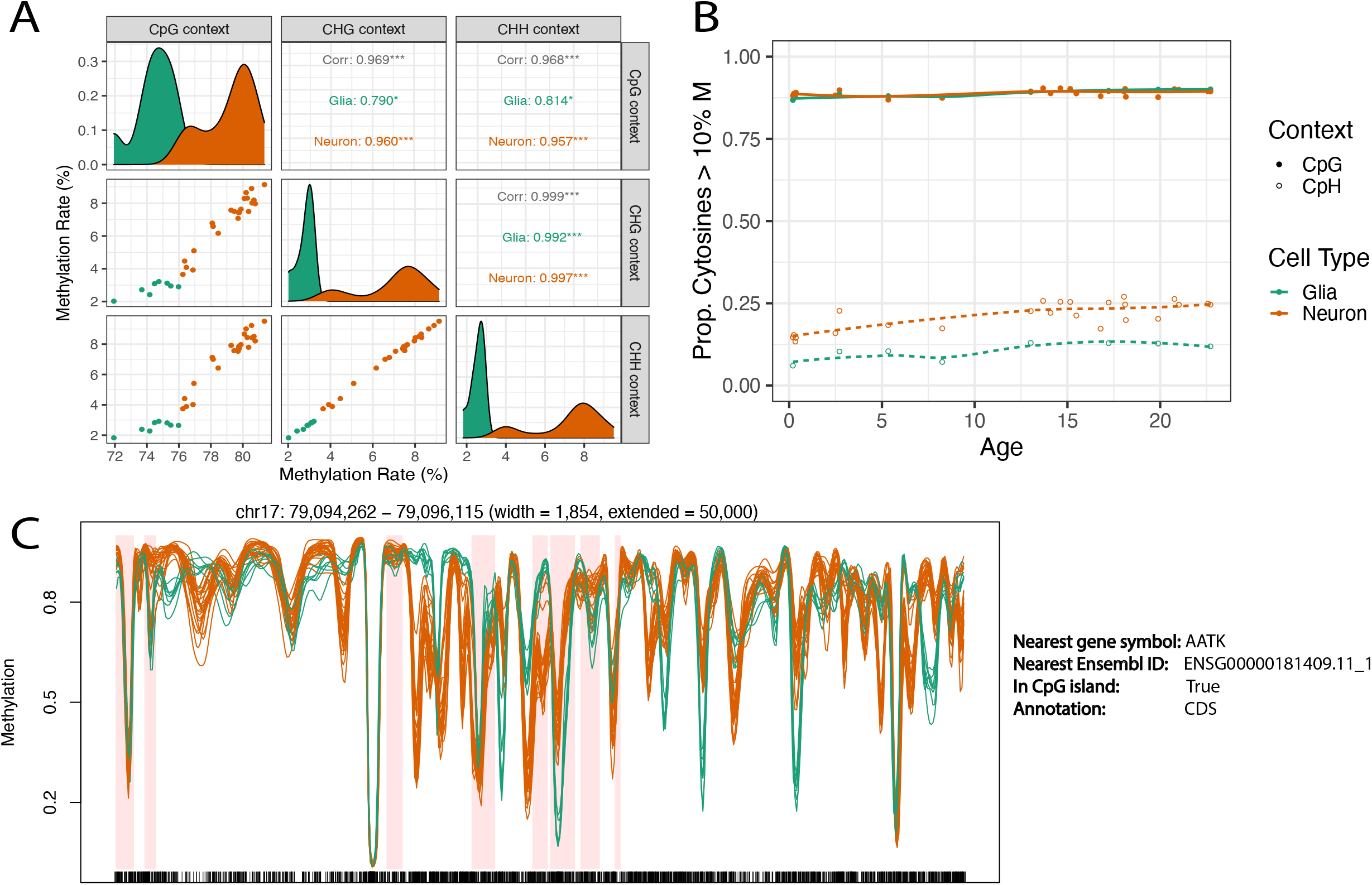
Visualization of BiocMAP outputs on an Example Dataset. **A** Comparison of average methylation rate by trinucleotide context and cell type, showing significant correlation between contexts and similar methylation distributions with different means by cell type. Glia have lower correlation with CpG and a lower methylation rate than neurons. **B** Proportion of highly methylated cytosines across age by cell type and trinucleotide context. Here it can be seen that neuronal CpH methylation appears to generally increase with age, with no obvious association with age in other combinations of cell type and context. **C** Genomic region containing differential methylation between neurons and glia. Orange and green methylation curves represent neuronal and glial samples, respectively. Windows highlighted in light red show differentially methylated regions (DMRs) determined in the Price et al. manuscript [5].

## Methods

### Overview

In the first alignment module, FASTQ files are checked with *FastQC* 0.11.8 [13], and by default samples that fail the “Adapter Content” metric are trimmed using *Trim Galore!* 0.6.6 [14] (see *Methods: Configuration* for alternative trimming options). The resulting FASTQ files are aligned to a reference genome using the GPU-based aligner *Arioc* 1.43 [7], to produce alignments in SAM format [17]. Using *SAMtools* 1.10 [17], alignments are filtered such that only primary alignments with *MAPQ* ≥ 5 are kept, duplicate reads are dropped using *SAMBLASTER* 0.1.26 [18], and finally the result is coordinate-sorted, indexed and stored in BAM format via *SAMtools*.

In the second extraction module, methylation extraction is performed with *MethylDackel* 0.5.2 [15] or optionally *Bismark* 0.23.0 [6] on alignments from the alignment module. By default, a BAM file and corresponding index are expected as input to *MethylDackel*, while just a SAM or BAM file is required if *Bismark* is to be used. For experiments using spike-ins of the lambda bacteriophage, we use a pseudo-alignment-based approach to infer bisulfite-conversion efficiency for each sample. In particular, we prepare two “versions” of the lambda genome: the original and a copy where each cytosine is replaced with a thymine (Availability of data and materials). The latter version represents “in-silico bisulfite conversion” of the original genome, where we assume this original is completely unmethylated. Using *kallisto* 0.46.1 [19], we count the number of reads aligned to each version of the lambda genome, and call these counts *o* and *b* for the original and bisulfite-converted versions, respectively. We define the bisulfite conversion efficiency rate e with the following ratio: 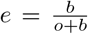. This contrasts with the more conventional approach [20], which involves directly aligning reads to the lambda reference genome and comparing cytosine and thymine counts on a single strand. In our own tests, we have found our pseudo-alignment-based approach to be sufficiently concordant with the conventional method, while requiring just a fraction of the computational time. The Bioconductor package *bsseq* [10] is used to gather methylation information into *SummarizedExperiment-based* [21] objects. The final result is a pair of *bsseq* objects, each of which contains all samples in the experiment as columns (**Figure 2**). One object contains all cytosines in the genome observed in CpG context, represented as rows, while the other contains the remaining cytosines in CpH context (**Figure 2**). Various metrics from *FastQC* [13], trimming, alignment, methylation extraction, and lambda pseudoalignment, if applicable, are aggregated into the *colData()* slot of each object (Additional File 1: S1).

### Configuration

A researcher typically must modify one or two types of files in BiocMAP: the execution script and optionally, a configuration file. The execution script contains the *nextflow* command [9] and major experiment-specific options, which together invoke one module in BiocMAP on a particular platform. Execution script templates are provided for each module in BiocMAP (“alignment” and “extraction”) and platform, which includes local Linux machines and systems managed by the Sun/ Son of Grid Engine (SGE) or Simple Linux Utility for Resource Management (SLURM) job schedulers (Additional File 1: S2). Additional job management systems are supported by Nextflow [9].

Required arguments in each execution script include “reference” and“sample”. “reference” may take values “hg38” or “hg19”, corresponding to the reference human genome to which samples should be aligned. The mouse reference genome “mm10” is also supported. “sample” may take values “single” or “paired”, referring to whether samples are single-end or paired-end. Several other options may be configured in the execution script. For example, the “input” argument takes the directory including *samples.manifest* for the alignment module, and the directory containing *rules.txt* for the extraction module (Additional File 1: S3). The optional “trim_mode” argument in the alignment module may take values “skip”, “adaptive”, or “force”, allowing the user to avoid trimming any samples, trim samples whose “Adapter Content” metric from *FastQC* [13] is “FAIL”, or trim all samples, respectively.

### Inputs

Both the alignment and extraction modules require a file called *samples.manifest* as input; this file is identical to those used in SPEAQeasy [22], thus providing a common file for describing input samples in both RNA-seq and WGBS projects. Briefly, *samples.manifest* is a tab-delimited plain-text file containing the absolute paths to each FASTQ file in the experiment, and associating files with a sample identifier. For compatibility with a previous format, optional MD5 sums can be associated with each FASTQ file. FASTQ files may optionally be compressed using gzip, so “.fq”, “.fastq”, “.fq.gz”, and “.fastq.gz” are all accepted file extensions. A researcher can specify that any combination of files be merged and treated as a single sample, simply by using the same sample ID in each line of FASTQ files to combine. This allows for simple management of the common case where a single biological sample is sequenced across several sequencing lanes and thus produces several files.

The extraction module makes use of the same *samples.manifest* file as the alignment module, as well as an additional file called *rules.txt* (Additional File 1: S3). The purpose of this file is to direct BiocMAP to each of the sets of outputs produced from the alignment module, which may not necessarily have been produced on the same high performance computing (HPC) system. From the alignment module, a researcher will have produced SAM or BAM files from alignment, as well as logs from quality checking, trimming, and alignment. The extraction module requires that a researcher create the *rules.txt* file, and point to the directory containing it via the *-input* option in the main submission script. *rules.txt* is a plain-text file where each line consists of a key-value pair (Additional File 1: S3). To facilitate this process, the alignment module creates a template *rules.txt* file which can be easily updated by the researcher in case file paths change in HPC system used for the extraction module. As an example *rule*, the “sam” key requires the path to each SAM or BAM from alignment as a value. Other keys are “manifest”, “arioc_log”, “xmc_log”, “trim_report”, “fastqc_log_first”, and “fastqc_log_last”, some of which are optional. Because an experiment typically consists of many samples, each value typically refers to multiple paths; including the literal “[id]” in a particular path signals for BiocMAP to replace “[id]” with each sample name to determine the full set of paths. A path can also be written as a glob expression, which is useful whenever a key refers to more than one file per sample; for example, the “sam” key can accept a BAM file and its index for every sample. For more detail about *rules.txt* and properly specifying file paths, see the documentation site (Availability of data and materials).

### Outputs

The final product from running BiocMAP is a pair of *bsseq* R objects, together containing methylation proportion and coverage information at all cytosine loci in the human or mouse genome (**Figure 2**). One object contains all cytosines occurring in “CpG” context in the reference genome, while the other object contains the remaining cytosines in “CpH” context. The *bsseq* [10] class extends the *SummarizedExperiment* [21] class, and here rows correspond to cytosines, while columns correspond to samples. Both objects contain the *M* and *Cov* assays, stored as *DelayedArray*s [23] and representing methylation proportions and coverage counts, respectively. We “strand-collapse” CpG loci, which involves combining methylation data from both genomic strands and thus discards strand-specific information. The CpG object is also smoothed using the *BSmooth* [24] pipeline as implemented in the *bsseq* [10] package, yielding an additional assay called *coef*, which contains the smoothing coefficients corresponding to each raw methylation proportion stored in the *M* assay. For a typical experiment involving many samples, these objects might occupy tens or even hundreds of gigabytes in memory if loaded in a traditional fashion. To enable working with the objects in a reasonable amount of memory, the assays are HDF5-backed using functionality from the *HDF5Array* [25] package. The HDF5 format is designed to allow direct manipulation of on-disk data as if it were loaded in random access memory (RAM), thus reducing the required RAM by an order of magnitude.

A table of metrics is stored in the *colData()* slot of each object, containing information collected from quality checking, trimming, alignment, methylation extraction, and potentially pseudoalignment to the lambda transcriptome (Additional File 1: S1). These metrics can be used for exploratory data analysis as well as for adjusting for them when performing downstream statistical analyses such as the identification of differentially methylated regions (DMRs). This table is also produced as a standalone R data frame to provide a format that is trivial to load into memory, interactively explore through https://libd.shinyapps.io/shinycsv/ (CITE https://zenodo.org/badge/latestdoi/72884509), or export to other formats.

In addition to the primary outputs of interest, BiocMAP produces a number of output files from intermediate pipeline steps (Additional File 1: S4).

### Software Management

BiocMAP is designed to run on Linux machines, either locally or through the SGE or SLURM job scheduling engines, or other engines supported by Nextflow [9]. We require Java 8 or later to be installed, as well as docker or singularity, based on the user’s preferred installation method. If neither are available, R [11] 4.0 or later and python 3 are required. Finally, an NVIDIA GPU and its corresponding drivers are required. Docker users must also have the NVIDIA container toolkit installed.

BiocMAP makes use of a number of external software tools which must be installed to use the pipeline. We support three different installation methods to accommodate a user’s existing set-up: download of docker images containing software, download of corresponding singularity images, or direct “local” installation of software. All three methods require simply invoking a shell script, followed by the name of the installation method (“docker”, “singularity”, or “local”):

~~~
bash install_software.sh “singularity”
~~~

For users of computing clusters, we make the assumption that GPU resources are accessible via a particular queue. Therefore, cluster users must also perform an additional manual step to complete the installation; this involves setting a variable in the appropriate configuration file (Additional File 1: S2) to the name of the queue where GPU(s) are available.

~~~
// The queue containing GPU access for use with Arioc
// (this must be set by the user!) arioc_queue =“gpu”
~~~

We recommend using the “docker” or “singularity” installation methods, if those tools are available, or the “local” method otherwise. As a Nextflow-based [9] pipeline, BiocMAP is out-of-the-box able to execute individual pipeline components, called *processes*, inside Docker or Singularity containers. These containers provide an exact environment, including the main software, system tools, and other dependencies, so that each BiocMAP process behaves identically on different computing systems. We host every docker image used by BiocMAP on a public Docker Hub repository (Availability of data and materials). In practice, the “singularity” installation mode automatically pulls the required docker images and builds singularity-compatible equivalents to use at run-time.

Alternatively, a researcher may use the “local” installation mode, which builds individual software tools from source when possible or downloads pre-compiled binaries otherwise. Since each piece of software is installed to a local directory and not globally, root permissions are not required for this installation method, and thus might be preferred by some users. However, because this approach tailors the installation to a particular computing environment, it is beyond our capacity to test unlike the “docker” or “singularity” modes, and we thus encourage you to avoid using the “local” mode.

### Annotation

Since BiocMAP performs alignment to a reference genome and can quantify lambda spike-ins [20], it must make use of external reference files. By default, required reference files are automatically pulled from GENCODE [26] (or NCBI for the lambda genome), but a user can also opt to provide their own files instead. The method used by BiocMAP to manage external reference files is nearly identical to that used in SPEAQeasy [22], and we encourage those interested to refer to that manuscript for more details; however, we provide a brief summary here.

When using default annotation from GENCODE [26], the genomes “hg38”, “hg19”, and “mm10” are supported; one of these values must be passed to the - -*reference* option. A researcher may specify the GENCODE version for human or mouse, as appropriate (e.g. “38” or “M27”, respectively). An additional configuration variable called “anno_build” determines if all sequences present in the “primary_assembly” file from GENCODE are kept, or if only canonical reference chromosomes are used for alignment; this corresponds to the values “primary” or “main” that a researcher may select, respectively. BiocMAP only pulls files from GENCODE that have not already been downloaded; after the first execution of the workflow for a given set of settings, it uses a locally cached copy of relevant files. A researcher may manually choose a directory to place annotation files via the command-line option ‘–annotation [path to directory]”, which enables potentially many users to share a single location for reference files to save disk space and time.

Alternatively, a researcher may provide their own reference genome in FASTA format in place of the automatically managed GENCODE [26] files. In this case, the “–annotation [path to directory]” option signifies the directory containing the provided FASTA file, and the “–custom_anno [label]” option assigns an informative label, or name, that can later be used in place of explicitly providing the genome. Note that the lambda genome is only automatically managed, since it is unlikely a user will need to swap out a different version.

### Test Samples

Small test files are provided in the *test* directory of the GitHub repository, for each combination of species (human and mouse) and pairing (single-end and paired-end). These are intended to allow a researcher to quickly verify BiocMAP has properly installed. While human, paired-end files are from the example *AgeNeunSortedWGBS* dataset [5, 16], the remaining files were retrieved from the Sequence Read Archive (SRA) (Additional File 1: S5). All FASTQ files were subsetted to 100,000 reads. A researcher can opt to run the extraction module on test data, without needing to run the alignment module beforehand. Test inputs to the extraction module, which include BAM files, their indices, and logs up through alignment, were generated by running BiocMAP with default settings, with the exception of using *trim.mode “force”* in place of the default *–trim_mode “adaptive”*.

### AgeNeunSortedWGBS Samples

The vignette provided with BiocMAP makes use of a dataset that includes 32 human postnatal dorsolateral prefrontal cortex samples up to 23 years of age [5, 16]. Homogenate postmortem tissue was sorted with NeuN-based fluorescence-activated nuclear sorting to produce 8 glial and 24 neuronal samples. The Price et al. manuscript also included prenatal samples that were excluded for this analysis [5].

*HDF5Array* 1.22.1 [25] is used to load the *bsseq* objects partially into memory, while keeping the *assays()* on disk as *DelayedMatrix* objects from the *DelayedArray*0.20.0 [23] package. Exploratory plots use *ggplot2* 3.3.5 [27] and the *ggpairs()* function from *GGally* 2.1.2 [28]. Methylation curves at a genomic region with several individual DMRs are explored with *plotRegion()* from *bsseq* 1.30.0 [10]. A total of 42GB of memory is required to run this analysis, and the analysis completes in 55 minutes.

## Discussion

Alignment to a reference genome is often the most computationally intensive component of a whole genome bisulfite sequencing (WGBS) data-processing workflow. As a result, workflows with an efficient alignment step can reduce total time required to process a dataset by a significant factor. CPU-based aligners like *Bismark* [6] or the more recent *BS-Seeker3* [29] can process WGBS samples in hours or days, but the GPU-based *Arioc* aligner offers higher alignment speeds than CPU-based aligners while maintaining comparable accuracy [7]. Thus we decided to support only *Arioc* in BiocMAP.

Given the recent introduction of GPUs and limited availability, researchers might not have access to GPUs on their main high performance computing (HPC) environment. HPC systems with GPUs might be under high demand or more expensive to use. Nextflow [9] does not provide functionality for executing some processing steps in one HPC system, transfering files, and resuming executing processes on a second HPC system. For these reasons, BiocMAP was implemented as two separate modules such that processing steps that benefit from the presence of GPUs can be run on HPC systems with GPUs, and the remaining steps can be run on regular CPU-powered HPC systems. Ultimately, if you have access to a HPC system with GPUs, you might prefer to run both modules on such a system. In that situation, BiocMAP’s two modules can be run serially without having to edit the *rules.txt* file that is automatically generated by the alignment module.

While BiocMAP is already likely to align reads quickly through *Arioc* with default settings, researchers are highly encouraged to configure BiocMAP settings to most efficiently use *Arioc* given the available hardware as noted on the documentation website (Availability of data and materials). Most configuration variables used by *Arioc* [7] can be directly edited in the appropriate BiocMAP configuration file (Additional File 1: S2). For example, the *batchSize* BiocMAP configuration variable is passed to the *batchSize* attribute of the *AriocU* or *AriocP* element of the configuration for *AriocU* or *AriocP*, which specifies how many reads *Arioc* can concurrently align per GPU utilized. This is one of many settings that depends on the specifications of the GPU(s) a researcher has available in their HPC system, which can be adjusted to achieve greater throughput. The *max_gpus* BiocMAP configuration variable specifies how many GPUs to use for alignment of each sample, potentially allowing increased parallelism when there is an abundance of GPU resources relative to number of samples in the experiment. A more comprehensive guide to adjusting BiocMAP configuration for a given computing environment is provided as part of the documentation website (Availability of data and materials).

We demonstrated how BiocMAP can be used to process publicly available WGBS data using an example dataset [5, 16]. Additional File 2 shows how you can then load the outputs of BiocMAP and use the data with R and Bioconductor packages such as *bsseq* [10], *ggplot2* [27], and *GGally* [28] to perform exploratory data analysis as well as downstream statistical analyses. In addition, development versions of BiocMAP were used in other peer reviewed publications [30, 31, 32] that have publicly available R code for several downstream analyses.

While we used a dataset of 32 samples to exemplify BiocMAP [5, 16], memory requirements scale roughly linearly with number of samples, with the production of a 600-sample dataset requiring about 200 GB of RAM, despite CpH and CpG cytosines encompassing around half of the genome (depending on the GC content of the genome; private WGBS datasets). Thus BiocMAP is scalable to a sample size larger than most if not all current WGBS datasets. The 200 GB RAM can likely be drastically reduced by future internal BiocMAP updates and can definitely be reduced for any dataset once you apply a filter on the number of reads per cytosine, stored in the *Cov* assay. Despite the potentially large memory requirements for running BiocMAP, loading the output *bsseq* [10] objects requires significantly less memory and is independent of the number of samples in the dataset, thanks to the HDF5 storage backend [25]. Users most likely need around 20-30 GB of RAM to load filtered *bsseq* objects for downstream statistical analyses for the CpH context, while the CpG context object requires less than 1 GB of RAM.

We envision that most users will not be interested in tweaking WGBS processing steps as long as they generate the output in a reasonable amount of time, but instead will want to focus on downstream analyses. We implemented BiocMAP in such a way that it will benefit from community developments in Bioconductor [12]. The main output data container is a *bsseq* [10] object that is an extension of *SummarizedExperiment* [21]. *SummarizedExperiment* itself is the one compatible with low-memory footprint backends such as *HDF5Array* [25]. If *SummarizedExperiment*becomes more efficient, by for example providing a low-memory footprint option for the gene coordinates (*rowRanges()* slot), users of BiocMAP will benefit from the reduction in RAM required to generate and load BiocMAP’s outputs. Similarly, if new R/Bioconductor packages are developed that implement downstream statistical analyses, they will be compatible with BiocMAP’s output objects as they are the central format for DNA methylation data [10, 12]. *zellkonverter* [33] is a Bioconductor package that allows exporting *SingleCellExperiment* R objects to Python. Given that *SingleCellExperiment* is an extension of *SummarizedExperiment*, just like *bsseq*, it seems reasonable to expect that *bsseq* objects will be readable from Python. Given these reasons, we envision that BiocMAP’s users will be able to use the resulting *bsseq* objects with any new methods implemented in R and most likely Python, two of the most widely used programming languages.

Despite the potential for customization within BiocMAP, it is designed to run “out of the box”, without a strict need to make hardware-specific configuration. This enables researchers to focus on their particular analysis questions instead of technical processing details.

## Conclusion

We implemented a whole genome bisulfite sequencing (WGBS) data processing workflow that relies on the GPU-accelerated *Arioc* aligner [7], yet is flexible enough to be used on multiple high performance computing (HPC) systems. The alignment output is further processed and packaged into *bsseq* [10] R/Bioconductor objects that are memory efficient and deeply integrated with the R/Bioconductor open source software ecosystem [12]. Thus BiocMAP will get the data processing job done in a fast and efficient manner for WGBS datasets up to several hundred samples, allowing researchers to focus their attention on exploratory data analysis and downstream statistical analyses. BiocMAP is available and documented at http://research.libd.org/BiocMAP/.

## Declarations

### Ethics approval and consent to participate

Not applicable.

### Consent for publication

Not applicable.

### Availability of data and materials

The BiocMAP software is available from GitHub at https://github.com/LieberInstitute/BiocMAP [8], with documentation at http://research.libd.org/BiocMAP/. The WGBS data used in the example vignette is available from https://www.synapse.org/#!Synapse:syn5842535 [5]. The original version of the lambda genome is available at ftp://ftp.ncbi.nlm.nih.gov/genomes/all/GCA/000/840/245/GCA_000840245.1_ViralProj14204/GCA_000840245.1_ViralProj14204_genomic.fna.gz. Docker images required for BiocMAP are hosted at https://hub.docker.com/orgs/libddocker/repositories.

### Author’s contributions

NJE and AEJ designed the software, then NJE and RW implemented the software. NJE and LCT wrote the manuscript draft. All authors contributed to revising the manuscript and approved the final manuscript.

## Acknowledgements

We would like to thank Anandita Rajpurohit and Kira A. Perzel Mandell (Lieber Institute for Brain Development), and Amanda J. Price (National Institute of Child Health and Human Development) for their suggestions on how to best present BiocMAP. Geo Pertea (LIBD) helped us proofread the documentation of BiocMAP.

## Authors’ information

Andrew E Jaffe is currently an employee and shareholder of Neumora Therapeutics, which is unrelated to the contents of this manuscript.

## Additional Files

Additional file 1 — Various tables with information about BiocMAP inputs, outputs, test files, and more.

This is provided as a multi-sheet excel file, with each sheet described in more detail below.

- S1: List of output metrics collected by BiocMAP. These are various quantities aggregated from processing steps like *FastQC* [13], trimming, alignment, and methylation extraction. Together they form an R *data.frame* accessible from the file *metrics.rda* and from within the *colData()* of output *bsseq* [10] objects. For paired-end samples, some metrics are computed separately for each mate, in which case metric names are appended with “_R1” and “_R2” to refer to each mate, respectively.
- S2: BiocMAP execution scripts and associated configuration files. BiocMAP provides several potential files for out-of-the-box functionality on local Linux machines as well as on SLURM or SGE-managed computing clusters.
- S3: Content of *rules.txt*. Each line of this input file to the extraction BiocMAP module consists of key-value pairs of the form < *key* >=< *value* >, some of which are required.
- S4: Intermediate output files. These files are not the main output files of interest from running both modules of BiocMAP, but are generated along the way as byproducts.
- S5: Sources of test data provided in the BiocMAP repository. Human and mouse single-end and paired-end samples are provided to allow users to quickly verify proper installation of BiocMAP, sourced from SRA or the FlowRNA-WGBS dataset.

Additional file 2 — Example vignette showing the use of BiocMAP output objects in downstream analysis.

This is a PDF file walking through R code and exploratory plots applied on the Price et al. data [16]. This file is also available from the BiocMAP GitHub repository (Availability of data and materials).

### Abbreviations

BiocMAP: **Bioc**onductor-friendly Nextflow-based **M**ethylation **A**nalysis **P**ipeline
CpG: Cytosine preceding a Guanine (methylation context)
CpH: Cytosine preceding a nucleotide other than Guanine (methylation context)
CHG: Cytosine followed by non-Guanine nucleotide, then Guanine (methylation context)
CHH: Cytosine followed by two nucleotides other than Guanine (methylation context)
CPU: Central Processing Unit
CUDA: Compute Unified Device Architecture
DMR: Differentially Methylated Region
GPU: Graphics Processing Unit
HPC: High Performance Computing
NCBI: National Center for Biotechnology Information
RAM: Random Access Memory
SGE: Sun Grid Engine or Son of Grid Engine
SLURM: Simple Linux Utility for Resource Management
SRA: Sequence Read Archive
WGBS: Whole Genome Bisulfite Sequencing

