## Additional File 2 for "BiocMAP: A Bioconductor-friendly, GPU-Accelerated Pipeline for Bisulfite-Sequencing Data"

### Example Analysis using BiocMAP Output Objects

**8 April 2022**

BiocMAP is a computational pipeline for bisulfite-sequencing data. Starting from the raw FASTQ-based sequencer outputs, BiocMAP ultimately produces two `bsseq` R objects, containing methylation proportions and coverage information for all genomic cytosines in CpG and CpH methylation context. Various metrics, such as bisulfite-conversion rate, number of reads trimmed, or concordant alignment rate, are collected and become columns in the `colData` of the `bsseq` objects. The below analysis will demonstrate how to explore both methylation information and quality metrics in the BiocMAP output objects. Background understanding of the bsseq package, and its parent SummarizedExperiment are helpful but not required here.

The dataset used here includes 32 human postnatal dorsolateral prefrontal cortex samples up to 23 years of age. Samples were derived using NeuN-based fluorescence-activated nuclear sorting. The associated manuscript is here.

### 1 Add Experiment Metadata to BiocMAP Outputs

The `bsseq` output objects from BiocMAP contain methylation and coverage info for our samples in the dataset. However, we're interested in exploring how this information relates back to sample metadata and phenotype information, present in an external file. Our first step will therefore be to load the BiocMAP output objects into memory, and manually attach the additional sample metadata to each object.

```
#  Load required R packages
library("bsseq")
library("HDF5Array")
library("ggplot2")
library("GGally")
library("tidyr")
library("here")
library("RColorBrewer")
```

```
#  Path to the sample metadata and BiocMAP outputs
meta_file <- here(
    "documentation", "example_analysis", "age_neun_pheno_data.csv"
)

#   Output bsseq objects from BiocMAP. A local directory is referenced because
#   the data is too large to host publicly
out_dir <- file.path(
    "/dcs04/lieber/lcolladotor/ageNeunSortedWGBS_LIBD001/ageNeunSortedWGBS",
    "processed-data/01-run_BiocMAP/pipeline_output/BSobjects/objects/combined"
)

#   CSV of DMRs already found for this dataset
dmr_list_path <- here(
    "documentation", "example_analysis", "age_neun_dmr_list.csv"
)
```

```
#  Load the 'CpG'-context object
bs_cpg <- loadHDF5SummarizedExperiment(dir = out_dir, prefix = "CpG")
```

```
#  Load the 'CpH'-context object. Note: this requires quite a bit of memory
#  (~23GB) even though the assays are disk-backed!
bs_cph <- loadHDF5SummarizedExperiment(dir = out_dir, prefix = "CpH")

#  Read in experiment-specific metadata and ensure sample ID orders match
meta <- read.csv(meta_file)
meta <- meta[match(colnames(bs_cpg), meta$Data.ID), ]

#  Add this metadata to the Bioconductor objects
colData(bs_cpg) <- cbind(colData(bs_cpg), meta)
colData(bs_cph) <- cbind(colData(bs_cph), meta)

#  Keep a copy of the metadata as a data frame, for easy plotting
meta_df <- data.frame(colData(bs_cpg))
```

We'll briefly look at how the data is distributed by age and cell type shown in Table 1 and Table 2, respectively.

```
cell_type_df <- data.frame(table(meta$Cell.Type))
colnames(cell_type_df) <- c("Cell Type", "Num Samples")
knitr::kable(
    cell_type_df,
    caption = "Distribution of cell type across samples",
    label = "cellTypes"
)
```

**Table 1:** Distribution of cell type across samples

| Cell Type | Num Samples |
|---|---:|
| Glia | 8 |
| Neuron | 24 |

```
age_bin_df <- data.frame(table(meta$Age.Bin))
colnames(age_bin_df) <- c("Age Bin", "Num Samples")
knitr::kable(
    age_bin_df,
    caption = "Distribution of age across samples",
    label = "ageBins"
)
```

**Table 2:** Distribution of age across samples

| Age Bin | Num Samples |
|---|---:|
| Child | 4 |
| Early.Teen | 5 |
| Neonate | 5 |
| Teen | 5 |
| Toddler | 3 |
| Young.Adult | 10 |

### 2 Exploratory Plots

#### 2.1 Bisulfite-Conversion Efficiency by Cell Type

This experiment used spike-ins of the lambda bacteriophage genome, which were quantified via BiocMAP to infer bisulfite-conversion rate. Successful bisulfite conversion is a pre-requisite for accurate methylation calls, so we'd like to see both that values (interpreted as percentages) are close to 100, and that values are not significantly different by sample (or by sample-related variables like cell type). We'll explore this visually in Figure 1.

```
ggplot(data = meta_df, aes(x = Cell.Type, y = lambda_bs_conv_eff)) +
    geom_boxplot(outlier.shape = NA) +
    geom_point() +
    labs(x = "Cell Type", y = "Bisulfite Conversion Rate (%)") +
    theme_bw(base_size = 20)
```

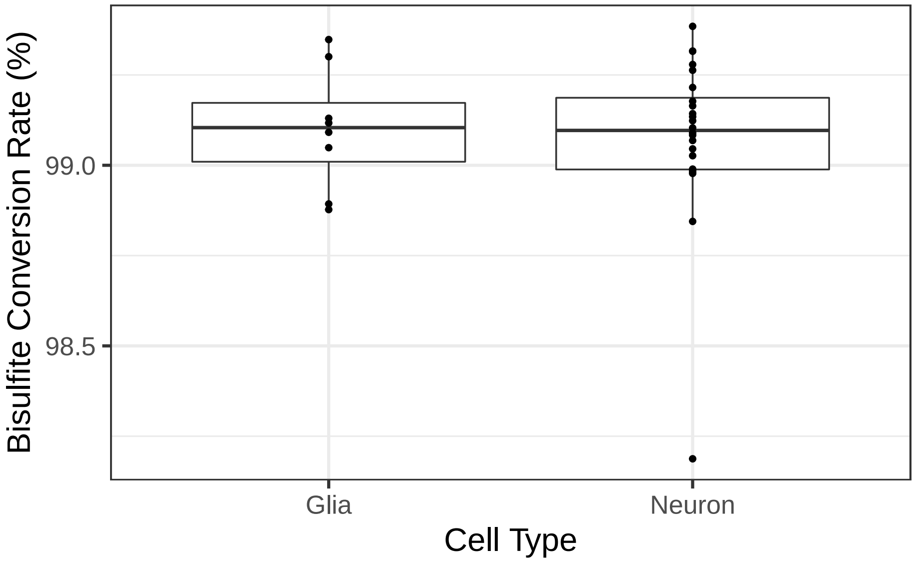

**Figure 1: Bisulfite conversion rate by cell type across samples**
Nearly all values are above 99% for both cell types, with one neuronal sample as low as just below 98.25%. Cell type does not appear to relate to bisulfite conversion rate, which is expected but reassuring.

#### 2.2 Relationship between Methylation Fractions across Cytosine Context, by Cell Type

Next, we'll explore if average methylation rate for each cytosine context correlates with that of other contexts across sample. For example, is a sample with highly methylated CpGs likely to have highly methylated CHGs? This is represented by the leftmost plot in Figure 2.

```
#   We'll make use of the 'ggpairs' function from the 'GGally' package, which is
#   well-suited for comparison of various metrics against each other, providing
#   density plots, comparison scatter plots, and correlation information.
```

```
ggpairs(
    data = meta_df,
    columns = c("perc_M_CpG", "perc_M_CHG", "perc_M_CHH"),
    xlab = "Methylation Rate (%)", ylab = "Methylation Rate (%)",
    columnLabels = c("CpG context", "CHG context", "CHH context"),
    mapping = aes(color = Cell.Type)
) +
    theme_bw(base_size = 15) +
    scale_color_brewer(palette = "Dark2") +
    scale_fill_brewer(palette = "Dark2")
```

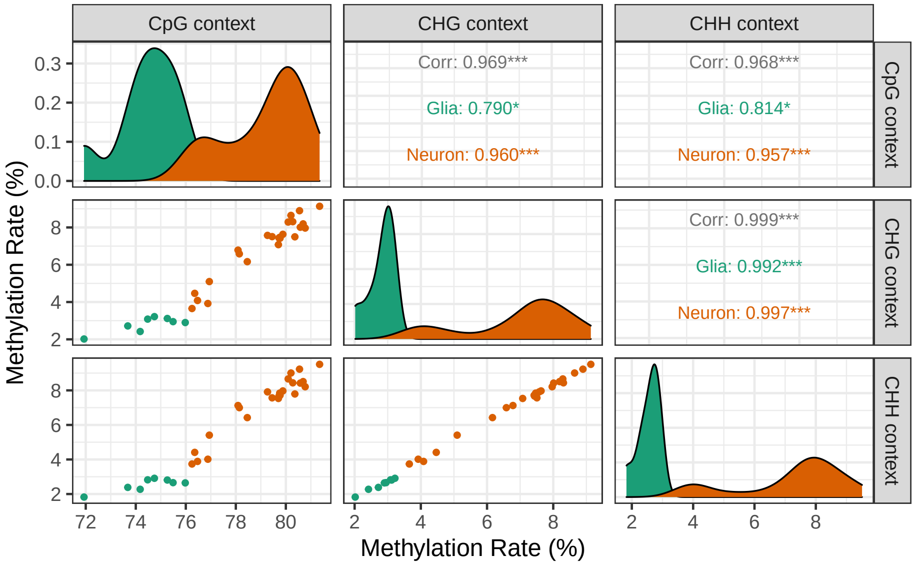

**Figure 2: Comparison of average methylation rate by cytosine context and cell type**
CpG and CpH methylation of both types strongly correlate with each other, with an apparent linear relationship in all cases. Density plots reveal that methylation distributions have approximately the same shape but different means within each cytosine context between cell types, though variances can differ within each CpH context. Finally a dramatic difference in CpH methylation can be seen between neuronal and glial samples, which is consistent with the literature; CpH methylation is known to be more prominent in neurons than many other cell types.

#### 2.3 Proportion of Highly Methylated Cytosines Across Age by Cell Type and Context

Another interesting area of exploration would be to examine how methylation patterns change with age. Note that we are exploring methylation for different donors of different ages, as the data does not include different observations across time for a fixed donor. In particular, we'll look at the proportion of cytosines for a given trinucleotide context that are at least 10% methylated (across all observations for a fixed sample).

```
#  The matrices in 'assays(bs_cpg)' and 'assays(bs_cph)' are stored on disk. To
#  speed up some below computations, we raise the per-block memory size
setAutoBlockSize(size = 1e9)
## automatic block size set to 1e+09 bytes (was 1e+08)
```

```r
#  Get the proportion of cytosines in each object (context) that have > 10%
#  methylation
meta_df$high_meth_cpg <- DelayedArray::colMeans(assays(bs_cpg)$M > 0.1)
meta_df$high_meth_cph <- DelayedArray::colMeans(assays(bs_cph)$M > 0.1)

#  Convert the data frame to "long" format for use with ggplot
meta_df_long <- meta_df %>%
    pivot_longer(
        cols = starts_with("high_meth_"),
        names_to = "context",
        names_prefix = "high_meth_",
        values_to = "high_meth_c"
    )
```

```r
ggplot(
    data = meta_df_long,
    aes(x = Age, y = high_meth_c, color = Cell.Type, shape = context)
) +
    geom_point() +
    geom_smooth(method = "loess", aes(linetype = context), se = FALSE) +
    scale_shape_manual(values = c(19, 1), labels = c("CpG", "CpH")) +
    scale_color_brewer(palette = "Dark2") +
    labs(
        x = "Age", y = "Prop. Cytosines > 10% M", color = "Cell Type",
        shape = "Context"
    ) +
    guides(linetype = "none") +
    theme_bw(base_size = 20) +
    ylim(0, 1)
## `geom_smooth()` using formula 'y ~ x'
```

#### 2.4 Explore a DMR

The next natural area of analysis is computing differentially methylated regions (DMRs) between groups of interest. In this dataset, we might explore methylation differences between neurons and glia. Since the corresponding manuscript already has done this, we'll read in the known DMRs from the supplementary table (additional file 2 here). Note that the `bsseq` package has an excellent guide for computing DMRs, a step that can be skipped in our case.

In this section, we'll read in the DMR list, "zoom out" around each DMR to form regions 100k bp wider (see `exploration_radius` below), and determine the region containing the most DMRs. We can then use `plotRegion` to visually check methylation levels at that region.

```r
exploration_radius <- 5e4

#   Read in CSV of DMRs and form a GenomicRanges object of unique regions
dmr_list <- read.csv(dmr_list_path)
dmr_gr <- GRanges(
    seqnames = dmr_list$Chromosome,
```

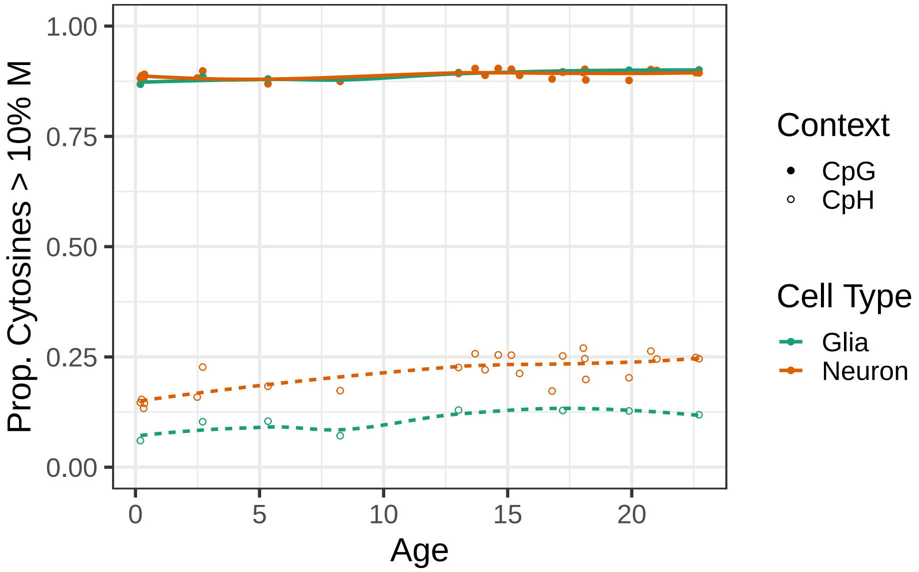

**Figure 3:** **Proportion of highly methylated cytosines across age by cell type and context**
While there isn't an apparent trend for CpG-context cytosines, CpH methylation show a clear increase with age, particularly for neurons.

```
        ranges = IRanges(dmr_list$Start, dmr_list$End)
    )
    dmr_gr <- unique(dmr_gr)

    #   Look around each DMR with a fixed radius
    dmr_gr_wide <- resize(
        dmr_gr,
        width = width(dmr_gr) + 2 * exploration_radius, fix = "center"
    )

    #   Find the DMR with the most nearby DMRs
    num_overlaps <- countOverlaps(dmr_gr, dmr_gr_wide)
    dmr_gr_overlap <- dmr_gr[match(max(num_overlaps), num_overlaps)]

    #   Color plots by cell type
    pal <- brewer.pal(3, "Dark2")[1:2]
    p_data <- pData(bs_cpg)
    p_data$col <- ifelse(bs_cpg$Cell.Type == "Glia", pal[1], pal[2])
    pData(bs_cpg) <- p_data
```

```
    plotRegion(
        BSseq = bs_cpg, region = dmr_gr_overlap,
        addRegions = dmr_gr, extend = exploration_radius
    )
```

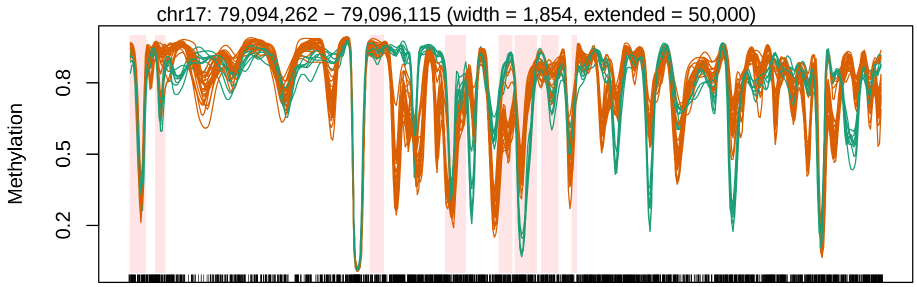

**Figure 4: Genomic region containing differential methylation between neurons and glia**
Orange methylation curves represent neuronal samples, whereas green curves represent glial samples. Windows highlighted in light red show DMRs determined in the manuscript between groups.

### 3 Conclusion

The above analysis was intended to show the nature of the BiocMAP output objects and touch on the many possibilities for exploratory data analysis and downstream statistical processing. We recommend the bsseq vignette for additional guidance to complement the analysis shown here. More generally, we hope BiocMAP is useful in connecting researchers to the vast analysis possibilities made possible by the hundreds of available Bioconductor packages.

### 4 Reproducibility Info

Date the vignette was generated:

```
## [1] "2022-04-08 17:44:47 EDT"
```

Wall-clock time spent generating the vignette:

```
## Time difference of 55.298 mins
```

Memory-related information while generating the vignette:

```
gc()
##              used    (Mb) gc trigger    (Mb)   max used    (Mb)
## Ncells    9444370   504.4   16671531   890.4   16671531   890.4
## Vcells 2933131405 22378.1 5430482561 41431.3 5430482561 41431.3
```

R session information:

```
## - Session info ----------------------------------------------------
##  setting  value
##  version  R version 4.1.2 Patched (2021-11-04 r81138)
##  os       CentOS Linux 7 (Core)
##  system   x86_64, linux-gnu
##  ui       X11
##  language (EN)
##  collate  en_US.UTF-8
##  ctype    en_US.UTF-8
##  tz       US/Eastern
```

```
## date     2022-04-08
## pandoc   2.13 @ /jhpce/shared/jhpce/core/conda/miniconda3-4.6.14/envs/svnR-4.1.x/bin/ (via rmarkdown)
##
## - Packages ---------------------------------------------------------
## package              * version  date (UTC) lib source
## assertthat             0.2.1    2019-03-21 [2] CRAN (R 4.1.0)
## Biobase              * 2.54.0   2021-10-26 [2] Bioconductor
## BiocGenerics         * 0.40.0   2021-10-26 [2] Bioconductor
## BiocIO                 1.4.0    2021-10-26 [2] Bioconductor
## BiocManager            1.30.16  2021-06-15 [2] CRAN (R 4.1.2)
## BiocParallel           1.28.3   2021-12-09 [2] Bioconductor
## BiocStyle            * 2.22.0   2021-10-26 [1] Bioconductor
## Biostrings             2.62.0   2021-10-26 [2] Bioconductor
## bitops                 1.0-7    2021-04-24 [2] CRAN (R 4.1.0)
## bookdown               0.24     2021-09-02 [1] CRAN (R 4.1.2)
## BSgenome               1.62.0   2021-10-26 [2] Bioconductor
## bsseq                * 1.30.0   2021-10-26 [2] Bioconductor
## cli                    3.2.0    2022-02-14 [2] CRAN (R 4.1.2)
## colorspace             2.0-3    2022-02-21 [2] CRAN (R 4.1.2)
## crayon                 1.5.1    2022-03-26 [2] CRAN (R 4.1.2)
## data.table             1.14.2   2021-09-27 [2] CRAN (R 4.1.2)
## DBI                    1.1.2    2021-12-20 [2] CRAN (R 4.1.2)
## DelayedArray         * 0.20.0   2021-10-26 [2] Bioconductor
## DelayedMatrixStats     1.16.0   2021-10-26 [2] Bioconductor
## digest                 0.6.29   2021-12-01 [2] CRAN (R 4.1.2)
## dplyr                  1.0.8    2022-02-08 [2] CRAN (R 4.1.2)
## ellipsis               0.3.2    2021-04-29 [2] CRAN (R 4.1.0)
## evaluate               0.15     2022-02-18 [2] CRAN (R 4.1.2)
## fansi                  1.0.3    2022-03-24 [2] CRAN (R 4.1.2)
## farver                 2.1.0    2021-02-28 [2] CRAN (R 4.1.0)
## fastmap                1.1.0    2021-01-25 [2] CRAN (R 4.1.0)
## generics               0.1.2    2022-01-31 [2] CRAN (R 4.1.2)
## GenomeInfoDb         * 1.30.1   2022-01-30 [2] Bioconductor
## GenomeInfoDbData       1.2.7    2021-11-01 [2] Bioconductor
## GenomicAlignments      1.30.0   2021-10-26 [2] Bioconductor
## GenomicRanges        * 1.46.1   2021-11-18 [2] Bioconductor
## GGally               * 2.1.2    2021-06-21 [2] CRAN (R 4.1.2)
## ggplot2              * 3.3.5    2021-06-25 [2] CRAN (R 4.1.0)
## glue                   1.6.2    2022-02-24 [2] CRAN (R 4.1.2)
## gtable                 0.3.0    2019-03-25 [2] CRAN (R 4.1.0)
## gtools                 3.9.2    2021-06-06 [2] CRAN (R 4.1.0)
## HDF5Array            * 1.22.1   2021-11-14 [2] Bioconductor
## here                 * 1.0.1    2020-12-13 [1] CRAN (R 4.1.1)
## htmltools              0.5.2    2021-08-25 [2] CRAN (R 4.1.2)
## httr                   1.4.2    2020-07-20 [2] CRAN (R 4.1.0)
## IRanges              * 2.28.0   2021-10-26 [2] Bioconductor
## jsonlite               1.8.0    2022-02-22 [2] CRAN (R 4.1.2)
## knitr                  1.38     2022-03-25 [2] CRAN (R 4.1.2)
## labeling               0.4.2    2020-10-20 [2] CRAN (R 4.1.0)
## lattice                0.20-45  2021-09-22 [3] CRAN (R 4.1.2)
## lifecycle              1.0.1    2021-09-24 [2] CRAN (R 4.1.2)
```

```
##  limma                    3.50.1   2022-02-17 [2] Bioconductor
##  locfit                   1.5-9.5  2022-03-03 [2] CRAN (R 4.1.2)
##  lubridate                1.8.0    2021-10-07 [2] CRAN (R 4.1.2)
##  magrittr                 2.0.3    2022-03-30 [2] CRAN (R 4.1.2)
##  Matrix                 * 1.4-1    2022-03-23 [3] CRAN (R 4.1.2)
##  MatrixGenerics         * 1.6.0    2021-10-26 [2] Bioconductor
##  matrixStats            * 0.61.0   2021-09-17 [2] CRAN (R 4.1.2)
##  mgcv                     1.8-40   2022-03-29 [3] CRAN (R 4.1.2)
##  munsell                  0.5.0    2018-06-12 [2] CRAN (R 4.1.0)
##  nlme                     3.1-157  2022-03-25 [3] CRAN (R 4.1.2)
##  permute                  0.9-7    2022-01-27 [2] CRAN (R 4.1.2)
##  pillar                   1.7.0    2022-02-01 [2] CRAN (R 4.1.2)
##  pkgconfig                2.0.3    2019-09-22 [2] CRAN (R 4.1.0)
##  plyr                     1.8.7    2022-03-24 [2] CRAN (R 4.1.2)
##  purrr                    0.3.4    2020-04-17 [2] CRAN (R 4.1.0)
##  R.methodsS3              1.8.1    2020-08-26 [2] CRAN (R 4.1.0)
##  R.oo                     1.24.0   2020-08-26 [2] CRAN (R 4.1.0)
##  R.utils                  2.11.0   2021-09-26 [2] CRAN (R 4.1.2)
##  R6                       2.5.1    2021-08-19 [2] CRAN (R 4.1.2)
##  RColorBrewer           * 1.1-3    2022-04-03 [2] CRAN (R 4.1.2)
##  Rcpp                     1.0.8.3  2022-03-17 [2] CRAN (R 4.1.2)
##  RCurl                    1.98-1.6 2022-02-08 [2] CRAN (R 4.1.2)
##  RefManageR             * 1.3.0    2020-11-13 [1] CRAN (R 4.1.2)
##  reshape                  0.8.8    2018-10-23 [2] CRAN (R 4.1.0)
##  restfulr                 0.0.13   2017-08-06 [2] CRAN (R 4.1.0)
##  rhdf5                  * 2.38.1   2022-03-10 [2] Bioconductor
##  rhdf5filters             1.6.0    2021-10-26 [2] Bioconductor
##  Rhdf5lib                 1.16.0   2021-10-26 [2] Bioconductor
##  rjson                    0.2.21   2022-01-09 [2] CRAN (R 4.1.2)
##  rlang                    1.0.2    2022-03-04 [2] CRAN (R 4.1.2)
##  rmarkdown                2.13     2022-03-10 [2] CRAN (R 4.1.2)
##  rprojroot                2.0.3    2022-04-02 [2] CRAN (R 4.1.2)
##  Rsamtools                2.10.0   2021-10-26 [2] Bioconductor
##  rtracklayer              1.54.0   2021-10-26 [2] Bioconductor
##  S4Vectors              * 0.32.4   2022-03-24 [2] Bioconductor
##  scales                   1.1.1    2020-05-11 [2] CRAN (R 4.1.0)
##  sessioninfo            * 1.2.2    2021-12-06 [2] CRAN (R 4.1.2)
##  sparseMatrixStats        1.6.0    2021-10-26 [2] Bioconductor
##  stringi                  1.7.6    2021-11-29 [2] CRAN (R 4.1.2)
##  stringr                  1.4.0    2019-02-10 [2] CRAN (R 4.1.0)
##  SummarizedExperiment   * 1.24.0   2021-10-26 [2] Bioconductor
##  tibble                   3.1.6    2021-11-07 [2] CRAN (R 4.1.2)
##  tidyr                  * 1.2.0    2022-02-01 [2] CRAN (R 4.1.2)
##  tidyselect               1.1.2    2022-02-21 [2] CRAN (R 4.1.2)
##  utf8                     1.2.2    2021-07-24 [2] CRAN (R 4.1.0)
##  vctrs                    0.4.0    2022-03-30 [2] CRAN (R 4.1.2)
##  withr                    2.5.0    2022-03-03 [2] CRAN (R 4.1.2)
##  xfun                     0.30     2022-03-02 [2] CRAN (R 4.1.2)
##  XML                      3.99-0.9 2022-02-24 [2] CRAN (R 4.1.2)
##  xml2                     1.3.3    2021-11-30 [2] CRAN (R 4.1.2)
##  XVector                  0.34.0   2021-10-26 [2] Bioconductor
```

```
## yaml                   2.3.5    2022-02-21 [2] CRAN (R 4.1.2)
## zlibbioc               1.40.0   2021-10-26 [2] Bioconductor
##
## [1] /users/neagles/R/4.1.x
## [2] /jhpce/shared/jhpce/core/conda/miniconda3-4.6.14/envs/svnR-4.1.x/R/4.1.x/lib64/R/site-library
## [3] /jhpce/shared/jhpce/core/conda/miniconda3-4.6.14/envs/svnR-4.1.x/R/4.1.x/lib64/R/library
##
## --------------------------------------------------------------------
```

### 5 Bibliography

This vignette was made possible by the following packages and software:

- R (R Core Team, 2021)
- *bsseq* = (Hansen, Langmead, and Irizarry, 2012)
- *BiocStyle* = (Oleś, 2021)
- *DelayedArray* = (Pagès, Hickey, and Lun, 2021)
- *GGally* = (Schloerke, Cook, Larmarange, Briatte, Marbach, Thoen, Elberg, and Crowley, 2021)
- *ggplot2* = (Wickham, 2016)
- *HDF5Array* = (Pagès, 2021)
- *here* = (Müller, 2020)
- *knitr* = (Xie, 2022)
- *RColorBrewer* = (Neuwirth, 2022)
- *RefManageR* = (McLean, 2017)
- *rmarkdown* = (Allaire, Xie, McPherson, Luraschi, Ushey, Atkins, Wickham, Cheng, Chang, and Iannone, 2022)
- *sessioninfo* = (Wickham, Chang, Flight, Müller, and Hester, 2021)
- *tidyr* = (Wickham and Girlich, 2022)
